# The role of CAPG in molecular communication between the embryo and the uterine endometrium: Is its function conserved in species with different implantation strategies?

**DOI:** 10.1101/2020.02.19.953794

**Authors:** Haidee Tinning, Alysha Taylor, Dapeng Wang, Bede Constantinides, Ruth Sutton, Georgios Oikonomou, Miguel A. Velazquez, Paul Thompson, Achim Treumann, Mary J O’Connell, Niamh Forde

## Abstract

During the pre-implantation period of pregnancy in eutherian mammals, changes to the uterine endometrium are required (both at the transcriptional and protein level) to facilitate the endometrium becoming receptive to an implanting embryo. We know that the developing conceptus (embryo and extraembryonic membranes) produces proteins during this developmental stage. We hypothesised that this common process in early pregnancy in eutheria may be facilitated by highly conserved conceptus-derived proteins such as macrophage capping protein CAPG. More specifically, we propose that CAPG may share functionality in modifying the transcriptome of the endometrial epithelial cells to facilitate receptivity to implantation in species with different implantation strategies, such as human and bovine. A recombinant bovine form of CAPG (91% sequence identity between bovine and human) was produced and bovine endometrial epithelial (bEECs) and stromal (bESCs) cells and human endometrial epithelial cells (hEECs) were cultured for 24 h with or without rbCAPG. RNA sequencing and quantitative real-time PCR analysis was used to assess the transcriptional response to rbCAPG (Control, vehicle, CAPG 10, 100, 1000 ng/ml: n=3 biological replicates per treatment per species). Treatment of bEECs with CAPG resulted in changes to 1052 transcripts (629 increased and 423 decreased) compared to vehicle controls, including those previously only identified as regulated by interferon-tau, the pregnancy recognition signal in cattle. Treatment of hEECs with bovine CAPG increased expression of transcripts previously known to interact with CAPG in different systems (*CAPZB, CAPZA2, ADD1* and *ADK*) compared with vehicle controls (P<0.05). In conclusion, we have demonstrated that CAPG, a highly conserved protein in eutherian mammals elicits a transcriptional response in the endometrial epithelium in two species with different implantation strategies that may facilitate uterine receptivity.

## INTRODUCTION

In most mammals, the majority of pregnancy loss occurs in the pre- and peri-implantation period of pregnancy. There are multiple factors such as poor gamete quality, and aberrant embryo development that can influence the likelihood of pregnancy loss. Selection processes of gametes and/or assisted reproductive technologies can overcome some of these issues. These advances/technologies have meant we can produce a large number of embryos *in vitro* that are viable to a specific stage of development (the blastocyst stage). These blastocysts can then be transferred into either patients (in humans) or into synchronised recipients (in domestic animals including cattle). However, pregnancy loss still occurs, thought to be due (in part at least) to inappropriate molecular communication between the developing conceptus (i.e. embryo and associated extraembryonic membranes) and the maternal environment – specifically the uterine endometrium (Lonergan and Forde, 2014; Macklon and Brosens, 2014).

The uterine endometrium is a complex tissue, consisting predominantly of luminal and glandular epithelial cells, and fibroblast-like stromal cells, alongside immune cells, and endothelial micro-vasculature. These heterogeneous cells are responsive spatially and temporally to steroid hormones - the concentrations of which fluctuate throughout the menstrual/estrous cycle in maternal circulation (Forde, Beltman, *et al.*, 2011; Bauersachs *et al.*, 2012). Following successful fertilisation, the embryo enters the uterus on day 4 of pregnancy where it is entirely reliant on the secretion and transport of molecules from the luminal and glandular epithelial cells of the endometrium. These molecules include growth factors, cytokines, amino acids and ions and are collectively termed uterine luminal fluid (ULF) (Mullen *et al.*, 2012; Forde *et al.*, 2014). ULF supports growth and development of the conceptus until a fully functional placenta is established (Burton *et al.*, 2002). The embryo undergoes a number of key morphological changes in the uterus including transition from a morula to a blastocyst where differentiation begins between the inner cell mass (that forms the foetus) and the outer cell mass (the trophectoderm, that goes on to form the placenta). Next, the blastocyst hatches from the zona pellucida, following which implantation occurs. However, implantation varies from species to species, for example in humans the blastocyst invades into the uterine endometrium (Carter, Enders and Pijnenborg, 2015) while in cattle the trophectoderm undergoes a rapid period of proliferation to produce an elongated conceptus and implantation does not begin to occur until Day 19 of pregnancy (Guillomot, Fléchon and Wintenberger-Torres, 1981). Irrespective of the early pregnancy morphology and type of implantation (epitheliocorial in human and cotyledonary in ungulates) two key conserved processes are required to modify the uterine endometrium to facilitate early pregnancy success, namely the actions of maternally-derived progesterone (P4) and the species-specific pregnancy recognition signal.

Firstly, P4 drives spatial and temporal changes to the endometrial transcriptome (Forde *et al.*, 2009). Paradoxically, continued exposure of the uterine endometrium to P4 down-regulates the P4 receptor from the luminal and glandular epithelium, a process required to establish uterine receptivity to implantation in all mammal species studied thus far (Kimmins and Maclaren, 2001; Okumu *et al.*, 2010). Secondly, the conceptus must secrete sufficient quantities of its pregnancy recognition signal to enhance uterine receptivity to implantation, i.e. modify the transcriptome and proteome of the endometrium to facilitate implantation as well as maintain P4 production. The molecular cue for pregnancy recognition varies across different mammals, e.g. in cattle and sheep it is a type 1 Interferon and in humans it is a chorionic gonadotrophin (hCG). However, the full extent of the essential interactions between the conceptus and the uterine environment required to support successful pregnancy have yet to be elucidated.

Previous studies by our group and others identified proteins produced by the peri-implantation conceptus in addition to the pregnancy recognition signal (Masters *et al.*, 1982; Forde *et al.*, 2015). We identified proteins uniquely present in the uterine fluid of pregnant cattle on the day of pregnancy recognition. These proteins were also detected in conceptus-conditioned medium with a large number of spectral counts via nano-liquid column-tandem mass spectrometry (Forde *et al.*, 2015). The protracted peri-implantation period of pregnancy in cattle provides opportunity to investigate the role of selected molecules in the process of pregnancy recognition. One of the proteins identified in the study by Forde *et al.*, (2015) is Macrophage Capping protein (CAPG) - a calcium-sensitive protein which functions to modulate cell motility through interactions with the cytoskeleton. CAPG therefore could function in remodelling the endometrium for successful implantation (Glaser *et al.*, 2014). CAPG also has known roles in cell migration, has been identified in exosomes from trophoblast cells *in vitro*, and has been shown to play a role in tumour invasiveness in endometrial cancer in humans (Martinez-Garcia *et al.*, 2017). Given its high abundance around maternal recognition of pregnancy in cattle (Forde *et al.*, 2015) and its roles in proliferation in human endometrial cells (Martinez-Garcia *et al.*, 2017) we hypothesised that CAPG modifies the endometrium during the pregnancy recognition process. This particularly important as the transcriptional response of the endometrium to the developing conceptus as a whole, is much greater than its response to the pregnancy recognition signal alone (Bauersachs *et al.*, 2012) and this contributes to pregnancy success. Because of the high levels of sequence conservation of this protein in mammals, we hypothesised that it may also modify the endometrial transcriptome in species with different implantation strategies to bovine, e.g. human. Our specific aims were to 1) assess levels of sequence conservation of CAPG in representative eutherian mammal species, 2) produce a recombinant form of bovine CAPG, and 3) test how the cells of the endometrium from different species respond to this rbCAPG *in vitro*.

## MATERIALS AND METHODS

### Analysis of sequence homology for CAPG

Homologs for CAPG were identified across 20 species including 15 placental mammals (Tarver *et al.*, 2016) and five outgroup species, and extracted from Ensembl (Yates *et al.*, 2016) (Table 1). A multiple sequence alignment was generated using MAFFTv7.3 (Katoh, 2002) for the 19 species with CAPG gene annotation (none available for microbat) and the resultant alignment was visualized in SeaView (Gouy, Guindon and Gascuel, 2010). The Percent Identity Matrices were generated using Clustal Omega (Sievers *et al.*, 2011).

**Table 1.**
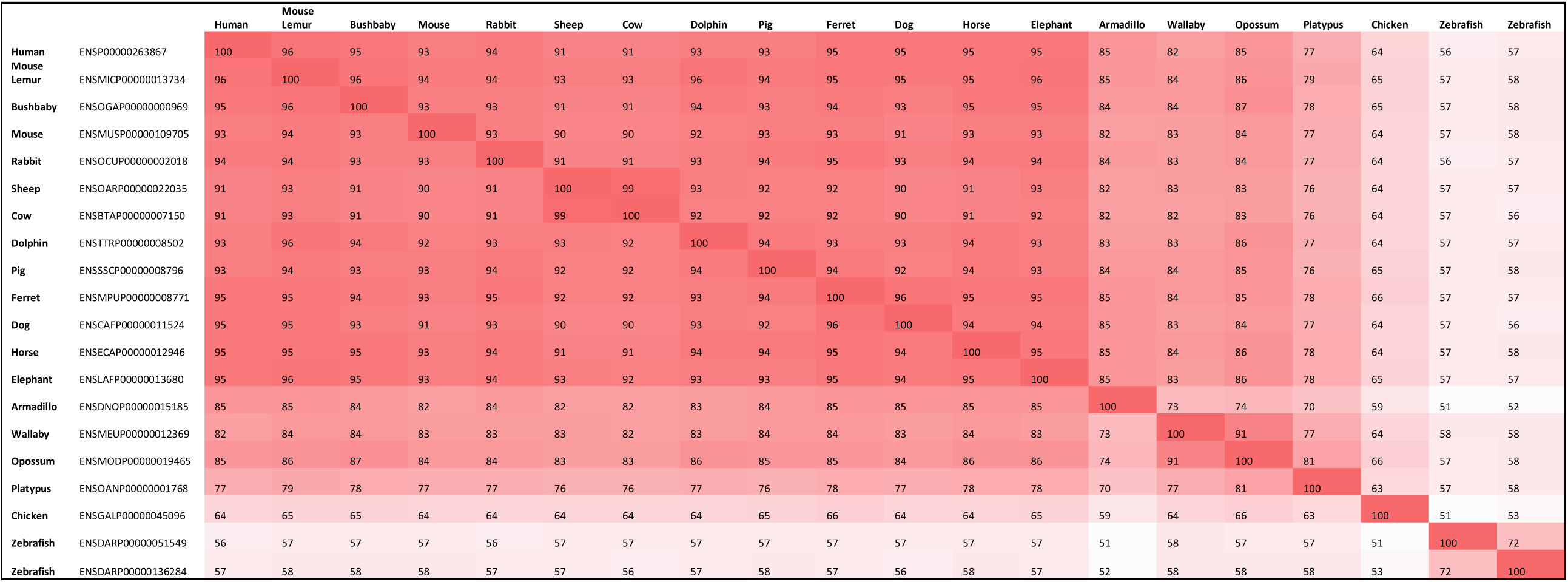
Levels of sequence identity for the CAPG homolog across a range of 20 species: 14 eutherian mammals, 2 marsupials, 1 monotreme, 2 birds, and 1 fish (2 orthologues). Darker cells indicate higher levels of sequence identity.

### Production of recombinant bovine CAPG (rbCAPG) protein

The sequence for bovine CAPG was subcloned into pET3A using ThermoFisher Scientific’s GenArt service, dissolved in 50 µl dH_2_O, and 2 µl of the plasmid transformed into HB101 pre-competent cells, selecting for transformants on 100 µl Ampicillin, Luria Agar plates. Two hundred ml Luria broth (100 µg/ml AMP) was inoculated with a scrape of plasmid, and were grown for 9 h at 37°C. The pellet was then harvested via centrifugation (9,000 rpm, 4°C, 10 min) and a midi prep was performed as per manufacturer’s instructions (Qiagen, Crawley, UK) and plasmids were dissolved in a final volume of 750 µl dH_2_O. Two µl of plasmid was transformed into BL21-AI pre-competent cells, selecting for transformants on 100µl Ampicillin, Luria Agar plates and grown overnight at 37°C. Twenty-four 10 ml LB starter cultures (100 µg/ml AMP), were inoculated with individual colonies picked from expression plates. These were then grown at 30°C for ∼ 24 h.

Large scale production of CAPG was undertaken by setup of 20 × 500ml LB (100 µg/ml AMP) flasks which were inoculated with the whole of an individual starter culture. These were held at 10°C until growth at 30°C, 160 rpm. All cultures were induced with a final concentration of 0.2% L-Arabinose 7 h later and cells were harvested cells by centrifugation (9,000 rpm, 4°C, 10 min), 5 h later. Each tube of cells was resuspended in 450 ml of 50 mM mixed phosphate, pH 7.2, 1 mM Benzamidine. The resulting cell suspension was sonicated at 14 microns for 42 min, whilst stirring on an ice bath, and then centrifuged at 9,000 rpm, for 42 min at 4°C. The supernatant was then carefully decanted and loaded onto a pre-equilibrated (Buffer 1) PROBOND column. The column was washed with 500 ml of Buffer 1, eluted with a 500 ml gradient of 0.0 M → 0.3 M Imidazole, 50 mM mixed phosphate, pH 7.2, and individual 5 ml fractions were collected.

Every third fraction from fraction 3-81 was subsequently loaded onto a 10% SDS-PAGE gel. Those fractions containing the largest/cleanest samples of CAPG were pooled and dialysed into 2 × 5 litre with 50mM mixed phosphate, pH7.2, along with 1mM DTT to remove Imidazole from the pool. Pooled CAPG was then filter sterilized with 0.45 µm disc filter and concentration/ quality calculated via UV spectroscopy.

### Bovine primary endometrial cell culture

Culture of bovine endometrial cells was achieved by isolating endometrial epithelial and stromal cells from late luteal stage bovine reproductive tracts as previously described (Ireland, Murphee and Coulson, 1980; Cronin *et al.*, 2012). Unless otherwise stated, all solutions were maintained at room temperature (RT). Individual tracts were sprayed with 70% EtOH and the ipsilateral uterine horn (the horn attached to the ovary with the corpus luteum (CL)) was opened longitudinally to expose the uterine endometrium. The endometrium was washed with 25 mL endometrial wash solution (DPBS, 1% ABAM) and dissected away from the underlying myometrium. The sheets of endometrium were placed into a wash solution (25 mL HBSS (no calcium no magnesium), 1% ABAM) and gently swirled prior to pour off and an additional 25 mL of HBSS was used to further wash endometrial strips. Endometrial tissue was then dissected into 3-5 mm^3^ pieces using tweezers and scissors and placed into 25 mL fresh HBSS. HBSS was poured off and digestive solution was added to the tissue to a final volume of 40 ml (Filter sterilised 50 mL HBSS (no calcium no magnesium), 25 mg collagenase II, 50 mg BSA, 125 uL 4% DNase I, 500 uL 0.0175% trypsin in HBSS) and tissue incubated in a rocking hot box for 1 h at 37°C.

The resulting solution was strained through a 100 µM strainer above a 40 µM strainer into a 50mL sterile falcon containing 5mL stop solution (HBSS with 10% charcoal-stripped FBS). The flow through, containing the stromal cell fraction, was vortexed briefly. The 40 µM strainer was inverted, flushed with 5 mL complete media (Gibco RPMI 1640, 10% charcoal-stripped FBS, 1% ABAM) (37°C) and the resulting epithelial-enriched fraction was plated into a T25 flask and cultured at 37°C in 5% CO_2_. Stromal cells were centrifuged at 400 g for 5 min and 5 mL of sterile H_2_O at 37°C was added to the pellet to lyse erythrocytes.

The stromal cell suspension was briefly vortexed and re-suspended immediately in 45 mL stop solution (37°C). The solution was then centrifuged at 400 g for 5 min, 10 mL of complete media (Gibco RPMI 1640, 10% charcoal-stripped FBS, 1% ABAM) (37°C) added to the pellet, re-suspended, and stromal cells were transferred into a T75 flask and placed into the 37°C/5% CO_2_ incubator. Twenty-four hours later (following adherence of cells) media was aspirated, washed 1 x with PBS (37°C) and 10 mL fresh media added. Once the stromal cells had reached ∼70% confluency, they were dislodged by trypsinisation (5 mL 0.025% trypsin in PBS) and plated at 150,000 per well into 6 well plates in 2 mL media for treatment.

Following 2-3 days of culture of epithelial cells at 37°C/5% CO_2_ un-adhered cells and media were aspirated and washed gently with 5 mL PBS (37°C) and replenished with 5 mL media. Contaminating stromal cells were removed by trypsinisation (0.025% trypsin in PBS) for 2 min. Cells were plated at 300,000 per well in 6-well dishes in 2 mL media, cultured at 37°C/5% CO_2_ overnight and media replenished prior to treatment.

Epithelial and stromal cells were then treated with one of the following for 24 h: 1) Control, 2) Vehicle, 3) 1000 ng/ml roIFNT, 4) 10 ng/ml rbCAPG, 5) 100 ng/ml rbCAPG, 6) 1000 ng/ml rbCAPG, 7) 1000 ng/ml rbCAPG+100 ng roIFNT. Following treatment, culture media was removed, cells were chemically lysed, snap-frozen in liquid nitrogen and kept at − 80°C.

### Human endometrial cell culture

Human immortalised endometrial epithelial Ishikawa cells (passage 10) were previously frozen in dimethyl sulfoxide (DMSO) and stored at −80°C. Cells were thawed rapidly, centrifuged at 400xg for 5 min, and re-suspended in 10 mL media (Dulbecco’s modified eagle medium: nutrient mixture F-12 (DMEM/F12), 10% foetal bovine serum (FBS), 1% GSP). Cells were then incubated in a T75 flask at 37°C/5% CO_2_, cultured until 80-90% confluent and split into T75 flasks (n=3). Human Ishikawa cells (n=3) were seeded at a density of 300,000 cells per well in 2 mL media, cultured at 37°C/5% CO_2_ overnight and media replenished prior to treatment. Cells were then treated with one of the following for 24 h: 1) Control, 2) Vehicle, 3) rbCAPG 10 ng/ml, 4) rbCAPG 100 ng/ml 5) rbCAPG 1000 ng/ml. Cells were chemically lysed, snap-frozen in liquid nitrogen and kept at −80°C.

### RNA extraction, library preparation and sequencing

Total RNA was extracted from cell lysate thawed on ice, using the MirVana RNA extraction kit as per manufacturer’s protocol (Life Technologies) and eluted in 50 µl RNase DNase free water. RNA was characterised using a Nanodrop and diluted to a standard concentration (Human 40 ng/ul; Bovine 50 ng/ul). Libraries were prepared from approximately 100ng total RNA in 10 µl of water using the Illumina TruSeq® Stranded Total kit, according to the manufacturer’s guidelines. rRNA was removed first by heating the RNA to 68°C for 5 min and cooling to room temperature for 1 minutes in the presence of 5 µl rRNA Removal Mix, followed by 1 min incubation at RT in the presence of 35 µl of rRNA Removal Beads. The beads were pelleted in the presence of a magnetic field and the solution containing mRNA and lncRNA removed and cleaned using Ampure RNAClean XP beads (Beckman Couter, California, USA) as follows: the RNA was bound to 99 µl of beads which were pelleted under magnetic field and the solution discarded. The beads were gently washed twice with 200 µl 70% ethanol and air dried. The beads were resuspended in 11 µl of Elution Buffer and 8.5 µl transferred to a new well to which 8.5 µl of the Elute, Prime, Fragment High Mix containing random hexamers was added prior to incubation at 94°C for 8 min. First strand synthesis was performed by the addition of 8 µl SuperScript II with First Strand Synthesis Act D Mix (1 to 9 ratio, respectively), and incubation at 25°C for 10 min followed by heating to 42°C for 15 min before terminating the reaction at 70°C for 15 min. Second strand cDNA synthesis was performed following the addition of 5 µl of resuspension buffer and 20 µl of Second Strand Master Mix containing DNA polymerase I and RNase H and incubation at 16°C for 1 h. Strand specificity is achieved by replacing dTTP with dUTP within the master mix. The cDNA was purified and size selected for fragments of approximately 300 bp by binding the cDNA to 90 µl AxyPrep Mag PCR clean up Kit (Axygen, Corning, New York, USA), the supernatant was discarded and the beads washed twice with 200 µl 80% ethanol, air dried and finally resuspended in 18.5 µl Resuspension Buffer. Following pelleting of the beads with a magnetic field, 17.5 µl of solution was transferred to a new well. Adenylation of 3’ ends was performed by adding 12.5 µl A-Tailing Mix and incubating the samples at 37°C for 30 minutes and terminated by heating to 70°C for 5 minutes, following the addition of 2.5 µl of Resuspension buffer. The samples were indexed by ligating 2.5 µl indexed Illumina sequencing compatible adapters to each sample in the presence of 2.5 µl Ligation Mix by heating of 30°C for 10 min. 5 µl of Stop Ligation Buffer was added to each well to stop the ligation process.

The samples were purified using 42 µl AxyPrep Mag PCR clean up Kit and washing twice with 80% ethanol and air-dried. The samples were resuspended in 52.5 µl Resuspension buffer and 50 µl transferred to a new well. The clean-up was repeated a second time with 50 µl of beads and resuspended in 22.5 µl of Resuspension buffer of which 20 µl was transferred to a new tube. The libraries were then amplified by PCR using 5 µl PCR Primer Cocktail and 25 µl PCR Master Mix with an initial 98°C for 30 sec, followed by 15 cycles of: 98°C 10 seconds, 60°C 30 sec, 72°C 30 sec with a final extension time of 72°C for 5 min. Finally, the cDNA libraries were purified and size selected to remove adaptor dimers and unincorporated adaptors using 50 µl AxyPrep Mag PCR clean up Kit as described above and resuspended in 32.5 µl Resuspension Buffer with 30 µl transferred to a new well.

The library quality, size range and sequencing adaptor dimer contamination was assessed with an Agilent TapeStation 2200 using the DNA broad range kit. Excess sequencing adaptor dimer if present was removed by AxyPrep Mag PCR clean up Kit bead mediated size selection. The final libraries were then quantified by measuring fluorescence with the Qubit dsDNA assay kit and Qubit fluorometer (Life Technologies) before creating an equimolar pool of the libraries. The RNA libraries were sequenced by the University of Leeds’s, Next Generation Sequencing Facility using the Illumina NextSeq 500 (Illumina, California, USA) with a single end 75 bp length read.

### RNA sequencing bioinformatics analysis and string interaction networks

To analyse differences in expression, adapter trimming was done by Cutadapt (Martin, 2011) and the reads were filtered using fastq_quality_filter as part of FASTX-toolkit with parameters including “-q 20” and “-p 90”. The reference genome and gene annotation files of cow (*Bos taurus*) and human species were retrieved from Ensembl genome database (release 96) (Cunningham *et al.*, 2019) and GENCODE (release 31) (Frankish *et al.*, 2019), respectively. With the help of Rsubread package (Liao, Smyth and Shi, 2019), read mapping was performed by means of Subread aligners with only uniquely mapped reads reported in the final alignment and read summarisation and quantification was carried out by featureCounts function. Statistical test for differential gene expression was conducted via DESeq2 (Love, Huber and Anders, 2014) with the cutoffs such as log2FoldChange > 1 (or < −1) and padj < 0.05. After this analysis, only protein-coding genes and lncRNAs were retained based on the gene biotype labels. Overrepresentation enrichment analysis of differential expression protein-coding gene sets was executed using WebGestalt (Liao *et al.*, 2019) for gene ontology terms and KEGG pathways, and protein-protein interaction networks were predicted using STRING (Szklarczyk *et al.*, 2019). For gene ontology terms, biological process non-redundant datasets were chosen as functional database, and for both types of analyses, significance level was determined by FDR < 0.05. For PCA plotting of each group of samples, both protein-coding genes and lncRNAs with RPKM value ≥ 1 in at least one sample were used and subsequently log2(RPKM+1) transformation and a quantile normalization were applied.

### Quantitative Real-time PCR analysis of selected transcripts in human and bovine endometrial cells

Transcripts analysed in bovine cells were selected on the basis of being upregulated during the pregnancy recognition process both due to and independent of IFNT. Selected transcripts for analysis in human cells were determined using string interaction network to identify nine human proteins that are known to be related to CAPG either through experimental determination or curated databases (Szklarczyk *et al.*, 2019). These were Cysteine Rich Protein 1 *(CRIP1)*, Ubiquitin-Fold Modifier Conjugating Enzyme 1 *(UFC1)*, Pirin *(PIR)*, Adenosine Kinase *(ADK)*, Ubiquitin-Fold Modifier 1 *(UFM1)*, Alpha-Adducin 1 *(ADD1)*, Capping Actin Protein of Muscle Z-Line Subunit Alpha 1 *(CAPZA1)*, Capping Actin Protein of Muscle Z-Line Subunit Alpha 2 *(CAPZA2)* and Capping Actin Protein of Muscle Z-Line Subunit Beta *(CAPZB).*

RNA was extracted as described above and RNA (500 ng bovine, 400 ng human) was reverse transcribed using the High Capacity cDNA reverse transcription kit (Applied Biosystems) as per manufacturer’s protocol. Nine human primers, seven bovine primers and respective normalisers (Table 2) were supplied by Integrated DNA Technologies (IDT) at 0.5 µM and 5 ng of cDNA per reaction was used alongside 5ul of 2x Roche SYBR green master mix and all samples were analysed in duplicate using the Roche Lightcycler 480 II. For human samples, *ACTB, GADPH* and *PPIA* were used as normaliser genes and for bovine samples, the average was calculated from *ACTB* and *GADPH* normaliser genes. The expression of genes of interest was determined using the comparative C_T_ method (2^-ΔΔCt^ method) (Livak and Schmittgen, 2001) prior to ANOVA to identify treatment effects on expression. The ANOVA analysis also included a Dunnett’s multiple comparisons test which allowed all sample data to be individually compared to the vehicle control data. Significant differences were determined when P <0.05.

**Table 2.**
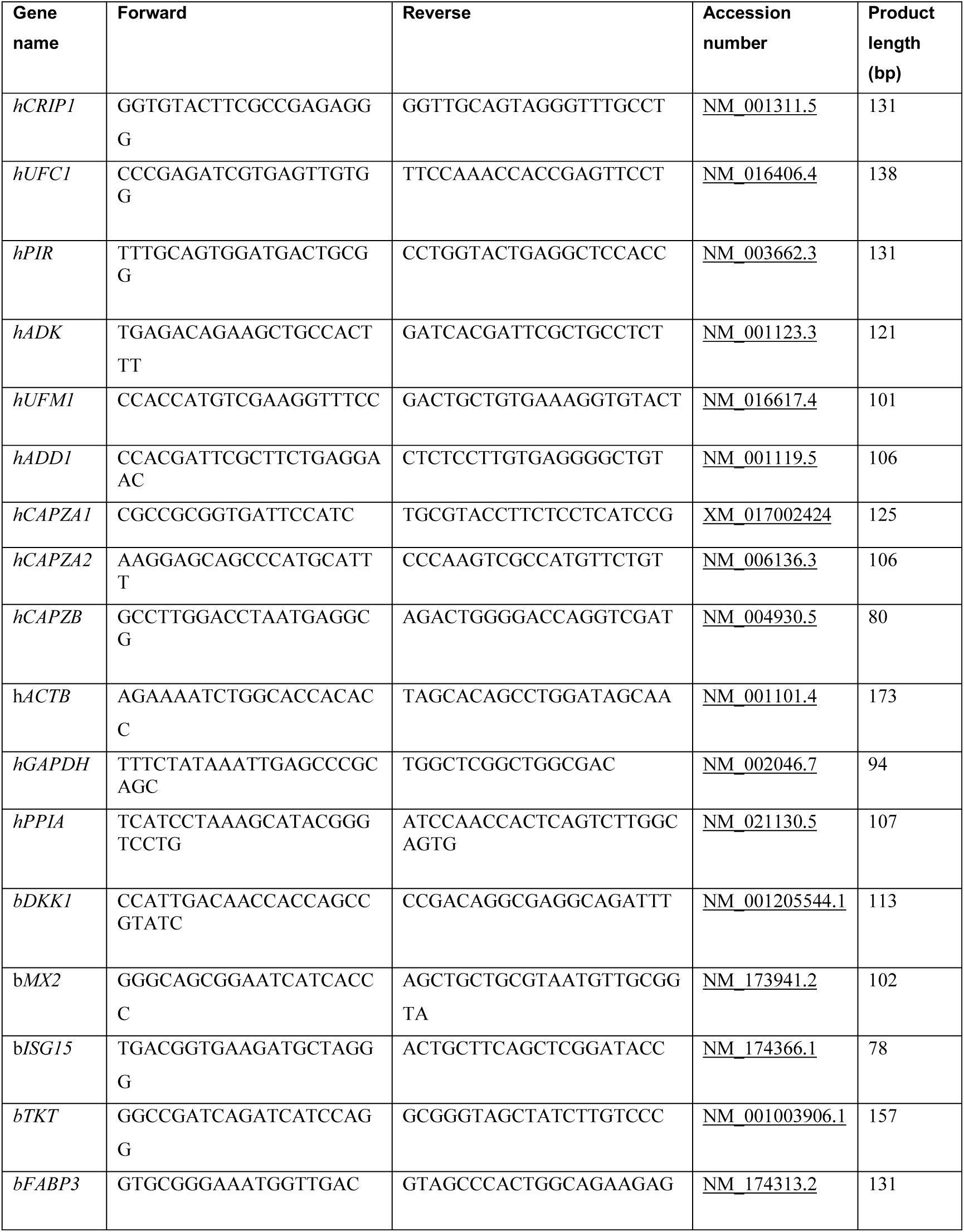
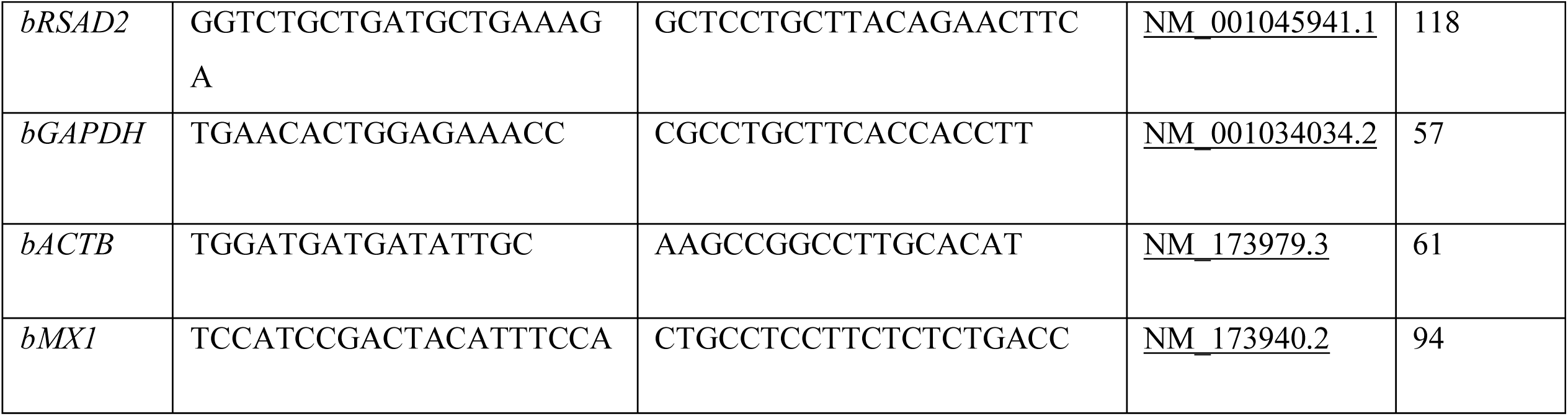
List of primers used for quantitative real-time PCR analysis of selected candidate transcripts in bovine and human endometrial cells treated with and without rbCAPG for 24 hours.

## RESULTS

### CAPG sequence is highly conserved across eutherian mammals

CAPG demonstrated a high degree of sequence conservation between all placental mammal species investigated (Table 1). Specifically, bovine CAPG showed > 90% sequence identity when compared to 12 of the 13 other placental mammals investigated, with the exception of the Armadillo CAPG gene (82% amino acid sequence identity). There was 99% similarity between bovine and ovine sequences, unsurprising given that these two species are closest relatives in the species sampled (diverging ∼18 MYA (Fernández and Vrba, 2005)). There was also a high degree of similarity compared to other placental mammals with different implantation strategies, for instance the human and bovine CAPG amino acid sequences are 91% identical.

### Treatment of bovine endometrial epithelial cells with rbCAPG induces a transcriptional response that may facilitate the pregnancy recognition process

Principal Component Analysis (PCA) of bovine endometrial epithelial cell gene expression data showed clear separation of control cells (no treatment) and those treated with +IFNT, +CAPG, or +IFNT&CAPG (Figure 1A). Treatment of epithelial cells for 24 hr with CAPG altered expression of 537 protein coding transcripts (422 increased and 115 decreased) compared to controls (Supplementary Table 1). One thousand four hundred and fifty-three transcripts were modified by IFNT alone (1028 increased and 452 decreased: Supplementary Table 2), while treatment of epithelial cells with a combination of CAPG and IFNT increased expression of 984 and decreased expression of 398 protein coding transcripts (Supplementary Table 3). Venn diagram analysis (Figure 2A) showed consistent changes in the expression of 400 transcripts when cells were treated with IFNT or CAPG alone or in combination (Supplementary Table 4). One hundred and twelve transcripts were modified with CAPG, and CAPG and IFNT, while 14 transcripts were modified by CAPG treatment alone. A string interaction network of differentially expressed transcripts in the epithelial cells, demonstrated that the majority of transcripts modified by IFNT alone, or in combination with CAPG, interacted with one another (Figures 3A and 3C). However, the string interaction network from CAPG treatment formed three distinct clusters of nodes indicating a difference in the transcriptional response induced by CAPG as compared to IFNT (Figure 3B).

**Figure 1.**
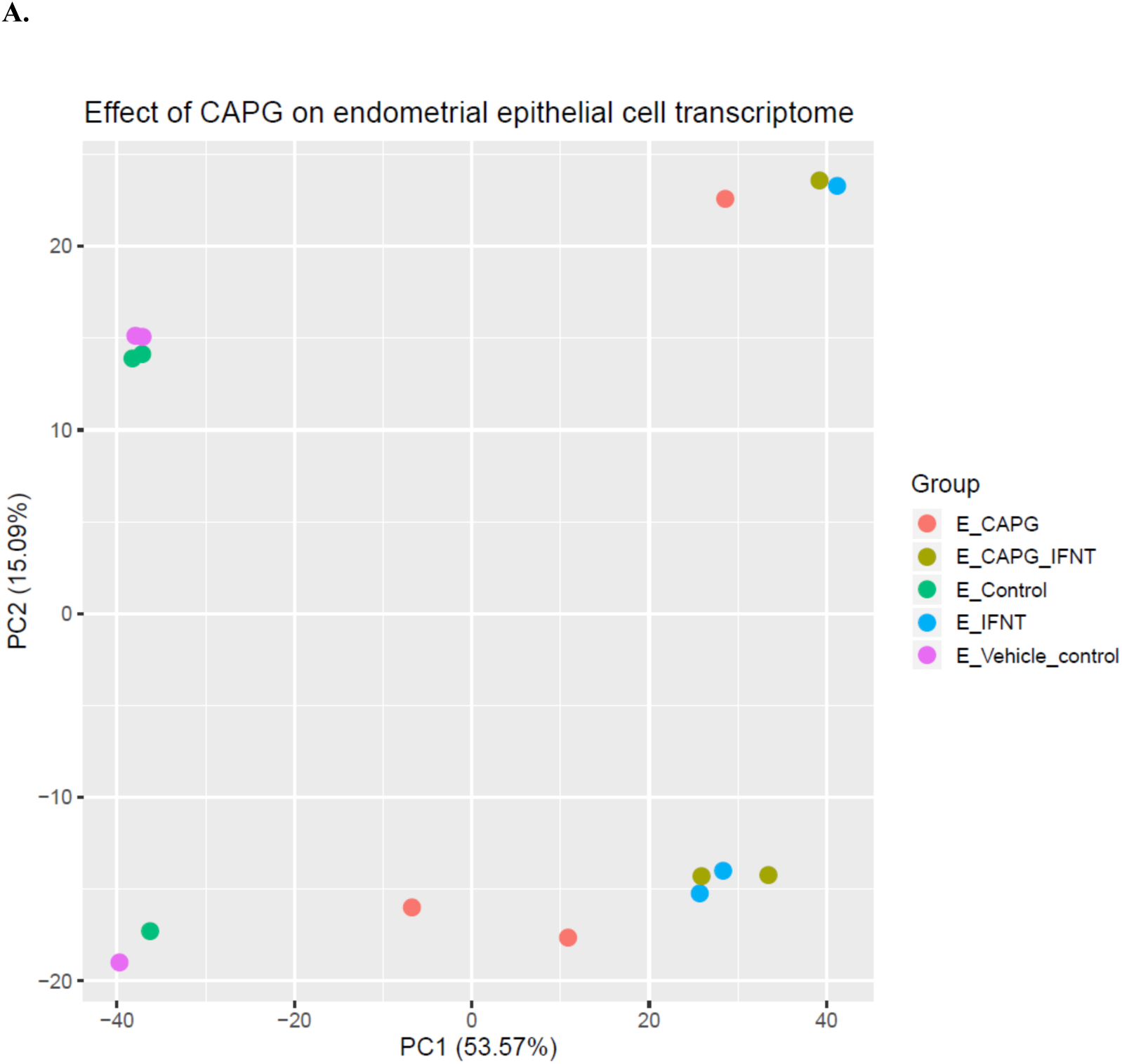
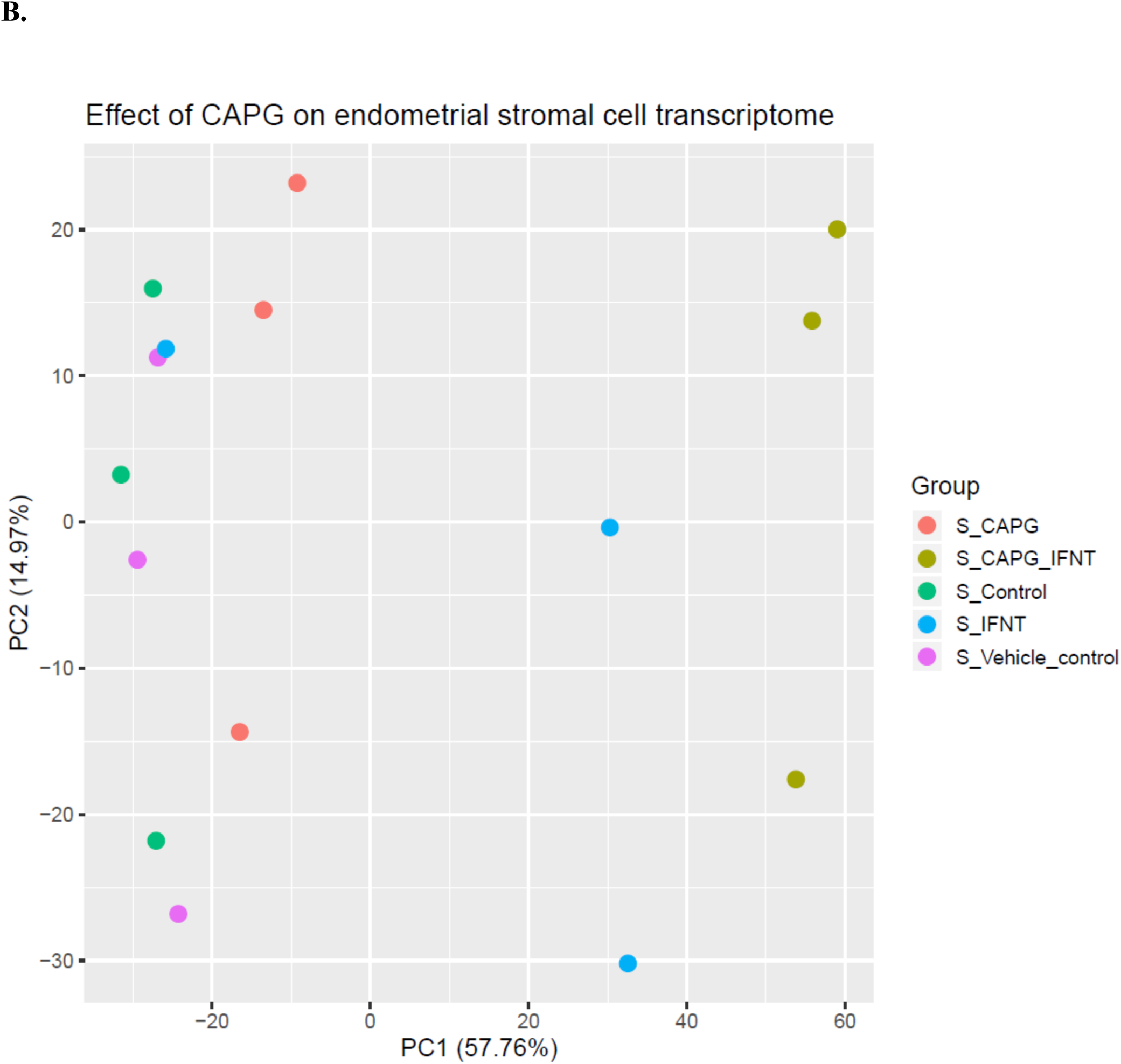
Principal component analysis (PCA) plot of overall transcriptional profile following RNA sequencing of A) bovine endometrial epithelial cells treated with 1000 ng/ml rbCAPG (E_CAPG), 1000 ng/ml rbCAPG + 100 ng/ml roIFNT (E_CAPG_IFNT), media control (E_Control), 1000 ng/ml of roIFNT (E_IFNT) or PBS vehicle control (E_Vehicle_control) for 24 hours or B) bovine endometrial stromal cells treated with 1000 ng/ml rbCAPG (S_CAPG), 1000 ng/ml rbCAPG + 100 ng/ml roIFNT (S_CAPG_IFNT), media control (S_Control), 1000 ng/ml of roIFNT (S_IFNT) or PBS vehicle control (S_Vehicle_control) for 24 hours.

**Figure 2.**
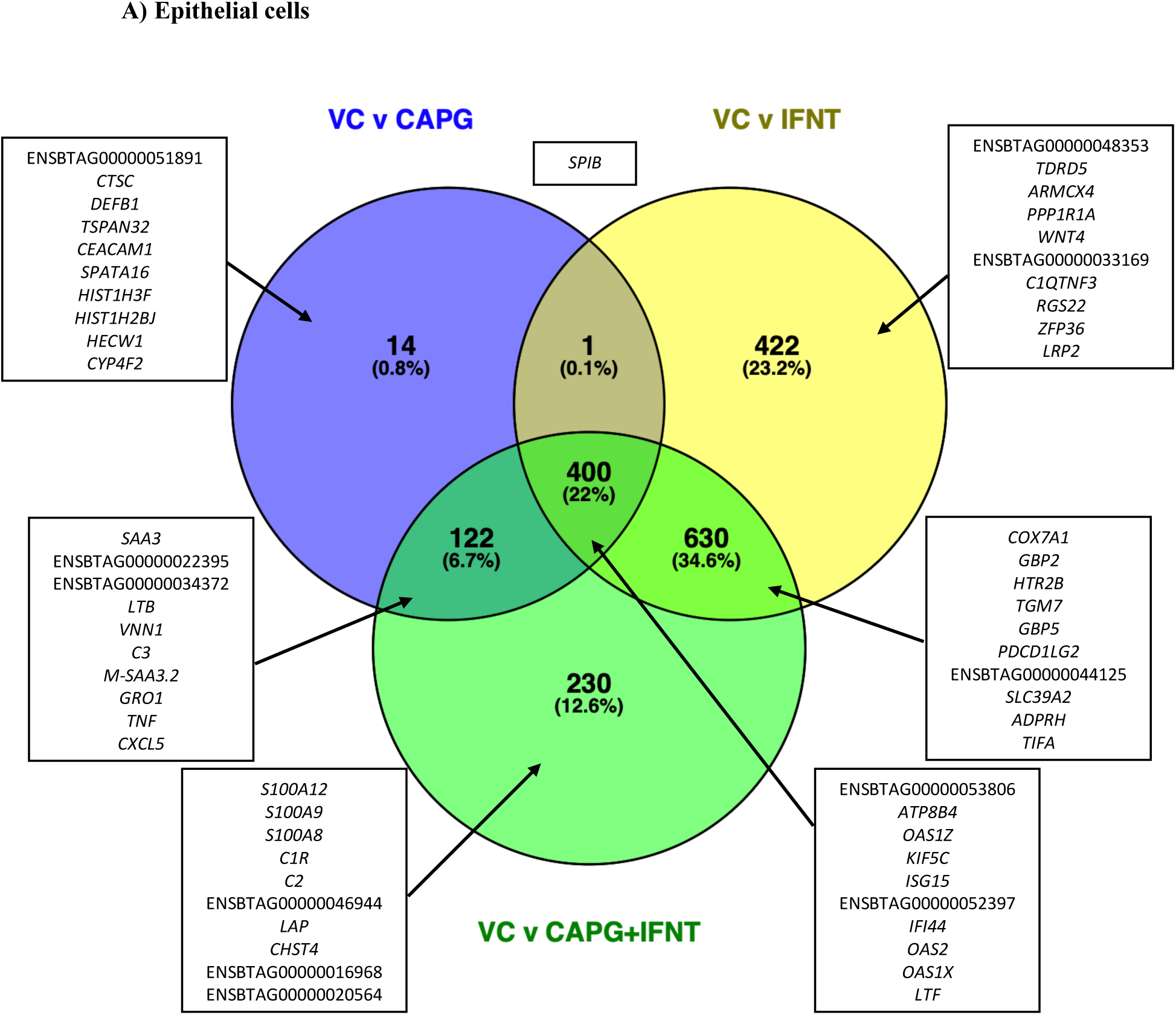
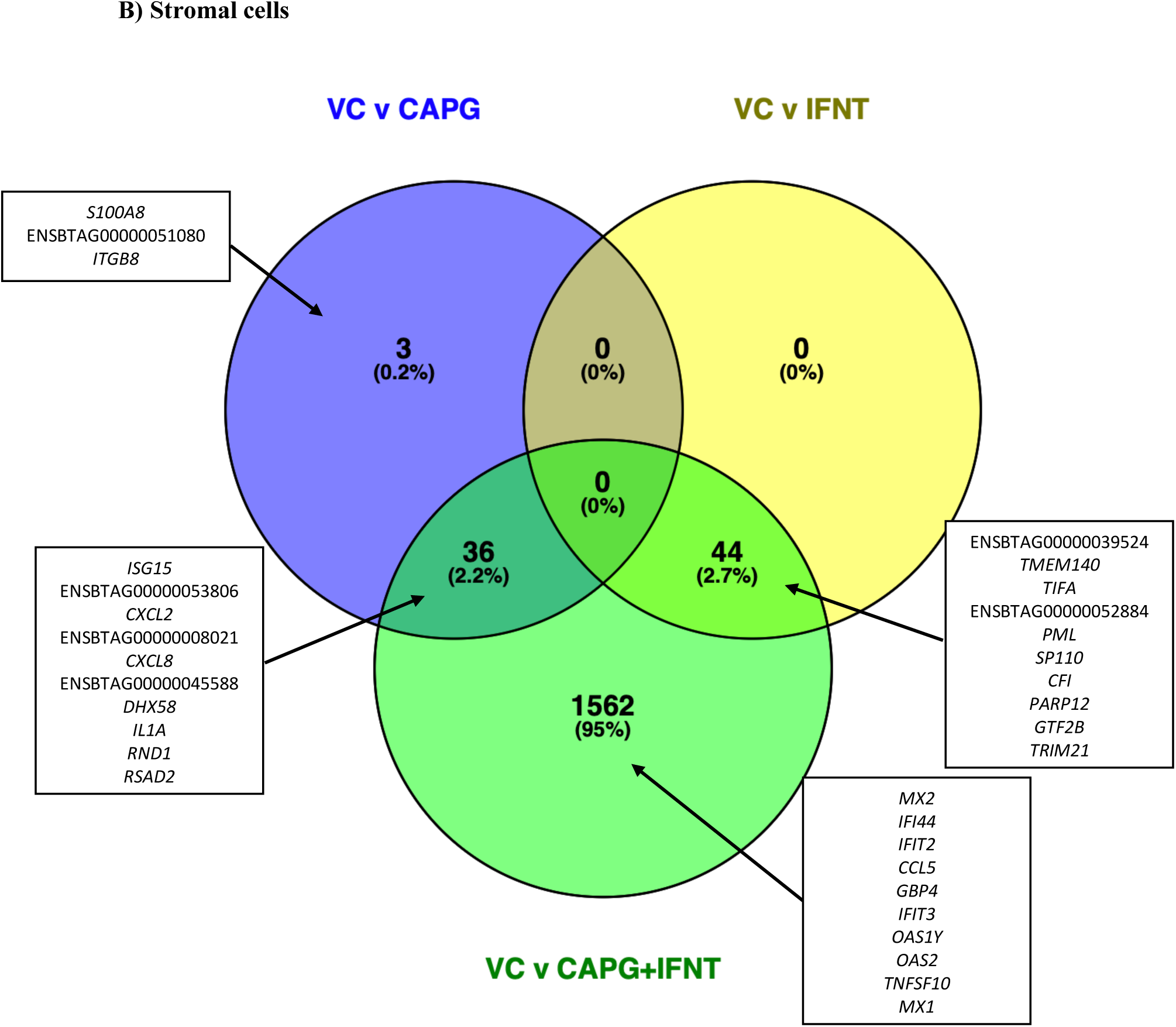
Venn diagram of differentially expressed transcripts with designated ensemble transcript identifiers with an adjusted p value of <0.05 and fold change of greater than two in **A)** bovine endometrial epithelial cells, or **B)** bovine endometrial stromal cells, treated with recombinant bovine CAPG (1000 ng/ml), recombinant ovine IFNT (1000 ng/ml) or a combination of both. All lists are comparisons between treatment and vehicle control and gene information is presented in Supplementary Tables 4 & 11. The top ten up/down regulated transcripts with the greatest fold change are shown in the boxes.

**Figure 3.**
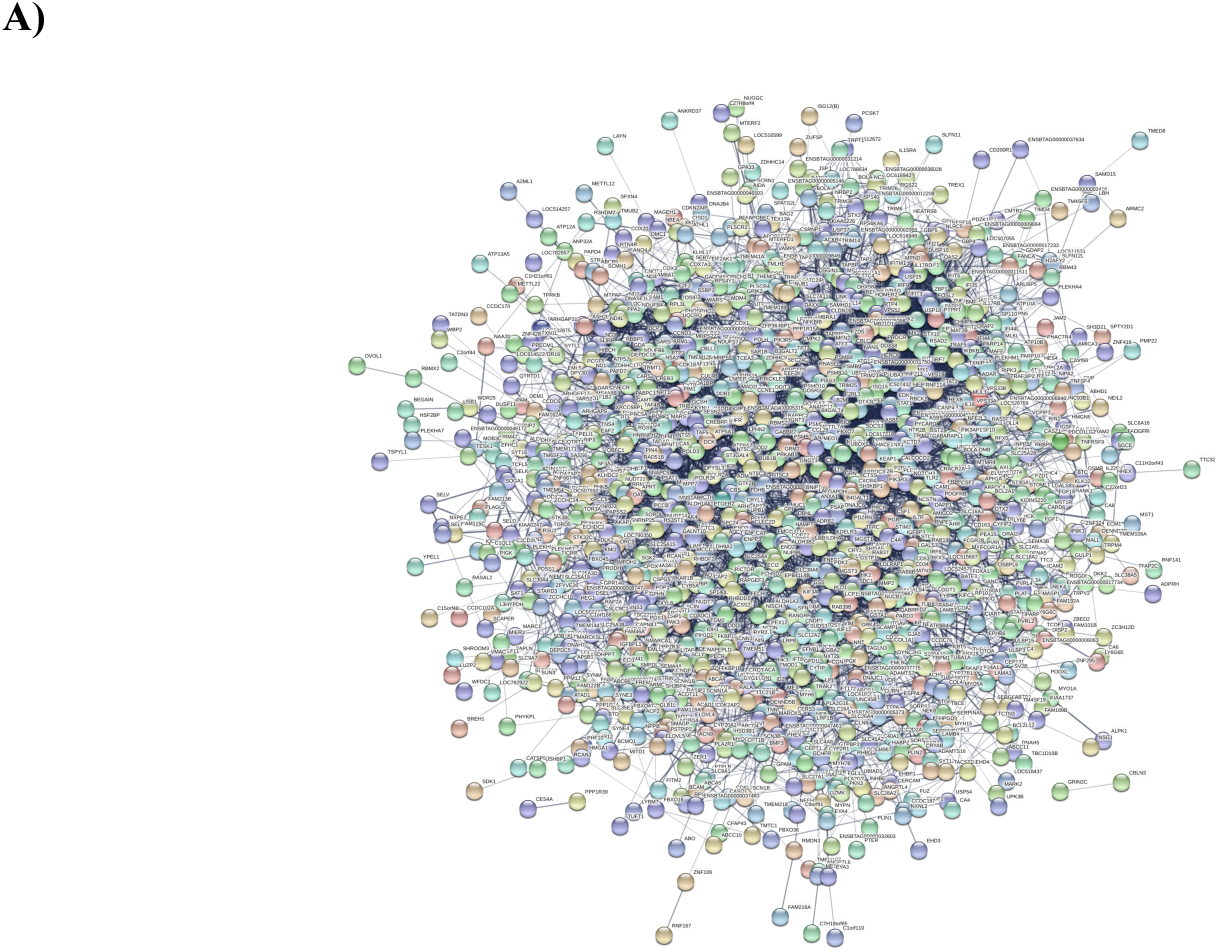
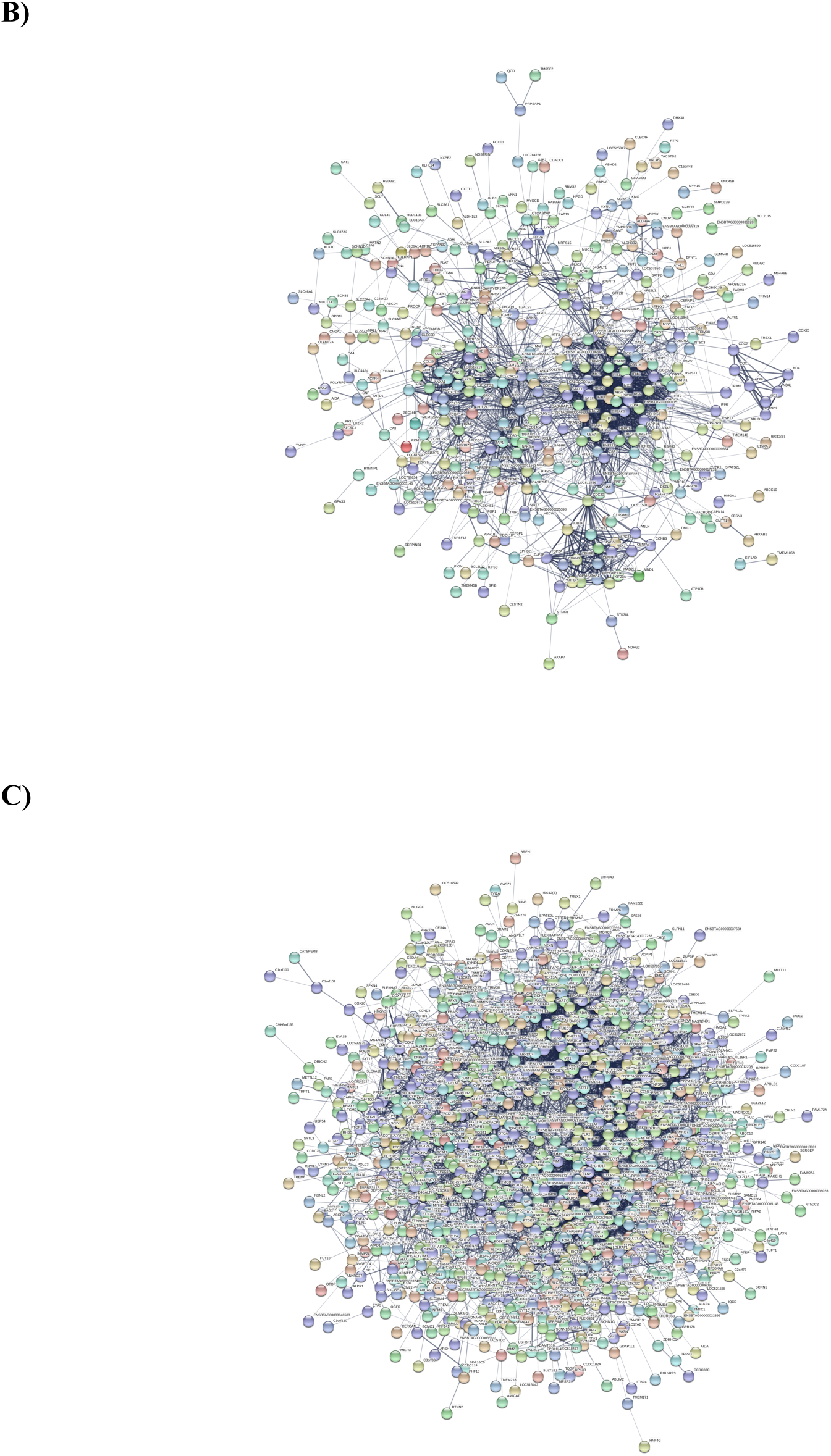
String interaction network analysis for those DEGs identified with an adjusted p value of <0.05 and with a fold change difference of fold change of greater than two following treatment of bovine endometrial epithelial cells for 24 hours with **A)** recombinant ovine IFNT (1000 ng/ml) alone, **B)** recombinant bovine CAPG alone (1000 ng/ml) alone or **C)** a combination of both for 24 hours.

### Specific gene ontologies and pathways were overrepresented following treatment of bovine endometrial epithelial cells with rbCAPG, roIFNT or a combination of both

Treatment of cells with CAPG for 24 h resulted in the modification of more transcripts involved in 29 biological processes than one would expect by chance (Supplementary Table 5). The GO term response to biotic stimulus contains 399 genes, 48 of which were altered by CAPG when we would have expected 12.5 DEGs to be changed by chance alone. Similarly, for GO terms response to cytokine 40 DEGs were overrepresented in our analysis from a possible 392 genes in this GO term, innate immune response (35/273), inflammatory response (32/292), immune effector process (32/301), positive regulation of immune system process (31/408), cytokine production (30/327), cellular response to oxygen-containing compound (28/434), regulation of defence response (27/256), regulation of response to external stimulus (27/325), regulation of immune response (26/315), nucleobase-containing small molecule metabolic process (26/384), small molecule biosynthetic process (25/372), cell activation (25/389), regulation of multi-organism process (23/179), drug metabolic process (23/384), response to lipid (22/335), import into cell (22/343), interspecies interaction between organisms (19/179), peptide secretion (19/255), purine-containing compound metabolic process (19/300), adaptive immune response (17/153), negative regulation of immune system process (17/190), leukocyte migration (14/160), response to oxidative stress (14/191), I-kappaB kinase/NF-kappaB signalling (13/118), antigen processing and presentation (9/40), cellular modified amino acid metabolic process (9/93), and humoral immune response (8/78).

In contrast, those that were regulated by IFNT treatment alone (Supplementary Table 6) involved 13 overrepresented biological processes including response to biotic stimulus (71/399), nucleobase-containing small molecule metabolic process (62/384), drug metabolic process (60/384), small molecule biosynthetic process (55/372), response to cytokine (53/392), innate immune response (48/273), purine-containing compound metabolic process (48/300), ribose phosphate metabolic process (45/281), immune effector process (45/301), organophosphate biosynthetic process (42/284), cofactor metabolic process (39/277), regulation of multi-organism process (29/179), and antigen processing and presentation (12/40).

### Specific pathways had significantly more transcripts modified following treatment of endometrial epithelial cells with rbCAPG

Similar to the overrepresented GO terms, more pathways were overrepresented following treatment with CAPG for 24 h than were expected by chance (Table 3). These included pathways associated with Epstein-Barr virus infection (24/177), NOD-like receptor signaling pathway (22/132), Influenza A (21/138), Cytokine-cytokine receptor interaction (20/227), Herpes simplex infection (19/147), Human immunodeficiency virus 1 infection (17/176), Kaposi sarcoma-associated herpesvirus infection (16/157), Measles (16/107), Chemokine signaling pathway (14/152), TNF signaling pathway (14/100), NF-kappa B signaling pathway (13/81), Phagosome (12/134), Hepatitis B (12/126), Necroptosis (12/120), Hepatitis C (11/102), Toll-like receptor signaling pathway (11/81), Rheumatoid arthritis (11/78), IL-17 signaling pathway (11/78), Legionellosis (11/50), C-type lectin receptor signaling pathway (10/90), RIG-I-like receptor signaling pathway (10/54), Cytosolic DNA-sensing pathway (10/45), Systemic lupus erythematosus (8/68), Pertussis (8/67), Antigen processing and presentation (8/54), Malaria (7/45), Arginine and proline metabolism (6/41), and Intestinal immune network for IgA production (6/42).

**Table 3.** Overrepresented pathways i.e. those with more differentially expressed transcripts associated with a gene list than one would expect by chance in primary bovine endometrial epithelial cells following treatment with recombinant bovine CAPG (1000 ng/ml) for 24 hours.

| Gene Set | Pathway | Numbers of genes in set | Number of DEGs in set | Enrichment Ratio | P Value | False discovery rate |
| --- | --- | --- | --- | --- | --- | --- |
| bta04621 | NOD-like receptor signaling pathway | 132 | 22 | 4.559468 | 1.50E-09 | 4.77E-07 |
| bta05169 | Epstein-Barr virus infection | 177 | 24 | 3.709398 | 1.85E-08 | 2.07E-06 |
| bta05164 | Influenza A | 138 | 21 | 4.162992 | 1.95E-08 | 2.07E-06 |
| bta05134 | Legionellosis | 50 | 11 | 6.018498 | 1.28E-06 | 6.99E-05 |
| bta05168 | Herpes simplex infection | 147 | 19 | 3.535914 | 1.30E-06 | 6.99E-05 |
| bta05162 | Measles | 107 | 16 | 4.090738 | 1.31E-06 | 6.99E-05 |
| bta04623 | Cytosolic DNA-sensing pathway | 45 | 10 | 6.079291 | 3.58E-06 | 1.63E-04 |
| bta04064 | NF-kappa B signaling pathway | 81 | 13 | 4.390599 | 6.00E-06 | 2.39E-04 |
| bta04668 | TNF signaling pathway | 100 | 14 | 3.829953 | 1.35E-05 | 4.79E-04 |
| bta04622 | RIG-I-like receptor signaling pathway | 54 | 10 | 5.066075 | 2.02E-05 | 6.43E-04 |
| bta04657 | IL-17 signaling pathway | 78 | 11 | 3.858011 | 1.08E-04 | 0.002881 |
| bta05323 | Rheumatoid arthritis | 78 | 11 | 3.858011 | 1.08E-04 | 0.002881 |
| bta04620 | Toll-like receptor signaling pathway | 81 | 11 | 3.715122 | 1.53E-04 | 0.003764 |
| bta05167 | Kaposi sarcoma-associated herpesvirus infection | 157 | 16 | 2.787955 | 1.76E-04 | 0.004018 |
| bta04060 | Cytokine-cytokine receptor interaction | 227 | 20 | 2.410291 | 2.09E-04 | 0.00425 |
| bta05170 | Human immunodeficiency virus 1 infection | 176 | 17 | 2.642419 | 2.13E-04 | 0.00425 |
| bta04612 | Antigen processing and presentation | 54 | 8 | 4.05286 | 6.82E-04 | 0.012792 |
| bta05144 | Malaria | 45 | 7 | 4.255503 | 0.001097 | 0.019317 |
| bta05160 | Hepatitis C | 102 | 11 | 2.950244 | 0.001151 | 0.019317 |
| bta04062 | Chemokine signaling pathway | 152 | 14 | 2.519706 | 0.001237 | 0.019723 |
| bta04217 | Necroptosis | 120 | 12 | 2.735681 | 0.001353 | 0.020552 |
| bta04625 | C-type lectin receptor signaling pathway | 90 | 10 | 3.039645 | 0.001528 | 0.022149 |
| bta05161 | Hepatitis B | 126 | 12 | 2.60541 | 0.002061 | 0.02859 |
| bta05133 | Pertussis | 67 | 8 | 3.266484 | 0.002845 | 0.037821 |
| bta05322 | Systemic lupus erythematosus | 68 | 8 | 3.218448 | 0.003126 | 0.039886 |
| bta00330 | Arginine and proline metabolism | 41 | 6 | 4.003435 | 0.003425 | 0.040792 |
| bta04145 | Phagosome | 134 | 12 | 2.449863 | 0.003453 | 0.040792 |
| bta04672 | Intestinal immune network for IgA production | 42 | 6 | 3.908115 | 0.003877 | 0.044171 |

Cells treated with IFNT for 24 h had a larger number of transcripts associated with the following pathways than one would expect by change (Supplementary Table 7), Metabolic pathways (135/1131), Herpes simplex infection (30/147), Epstein-Barr virus infection (29/177), Influenza A (27/138), NOD-like receptor signaling pathway (26/132), Parkinson disease (24/132), Cytosolic DNA-sensing pathway (12/45), Arginine and proline metabolism (12/41), and Nicotinate and nicotinamide metabolism (10/25).

### Limited transcriptional response of bovine endometrial stromal cells to rbCAPG following 24 h of treatment

PCA analysis of stromal cells gene expression data demonstrated less clear separation between controls and treated cells compared with that observed in epithelial cells (Figure 1B). This was reflected in the number of proteins coding transcripts that were modified with only 38 increased in expression while 1 decreased following treatment with CAPG for 24 h (Supplementary Table 8) while treatment with IFNT modified 44 transcripts all of which were increased in expression (Supplementary Table 9). However, treatment of stromal cells with a combination of CAPG & IFNT modified the expression of 1642 transcripts, of which 1095 increased and 547 decreased in expression (Supplementary Table 10). Venn diagram analysis (Figure 2B) showed that only three transcripts were modified by CAPG alone (*S100A6, ITGB8* and an uncharacterised transcript) while all those transcripts modified by IFNT were also changed by the combined treatment of CAPG and IFNT (Supplementary Table 11). Given the limited number of DEGs identified in CAPG and IFNT alone treatment, no downstream analysis was undertaken.

### Selected transcripts are modified by rbCAPG in a dose-dependent manner in both human and bovine cells in vitro

RNA sequencing analyses of Ishikawa cells treated with 1000 ng/ml of bovine rbCAPG did not identify any differences in the transcriptional response (data not shown). However, qRT-PCR analysis of human endometrial epithelial cells exposed to rbCAPG in a dose-dependent manner, resulted in increased expression of *ADD1, ADDK, CAPZA2, CAPZB*, and *CRIP1*, but only at the lower doses of rbCAPG compared to vehicle control (P<0.05: Figure 5A) except in CAPZB. In bovine stromal cells, the expression of *MX1, MX2*, and *RSAD2* increased in a dose responsive manner but only when treated with rbCAPG in combination with IFNT (Figure 5B&C) while expression of *ISG15* and *TKT* had a limited response to rbCAPG treatment alone whereas the addition of IFNT and rbCAPG enhanced their expression (P < 0.05).

**Figure 4.**
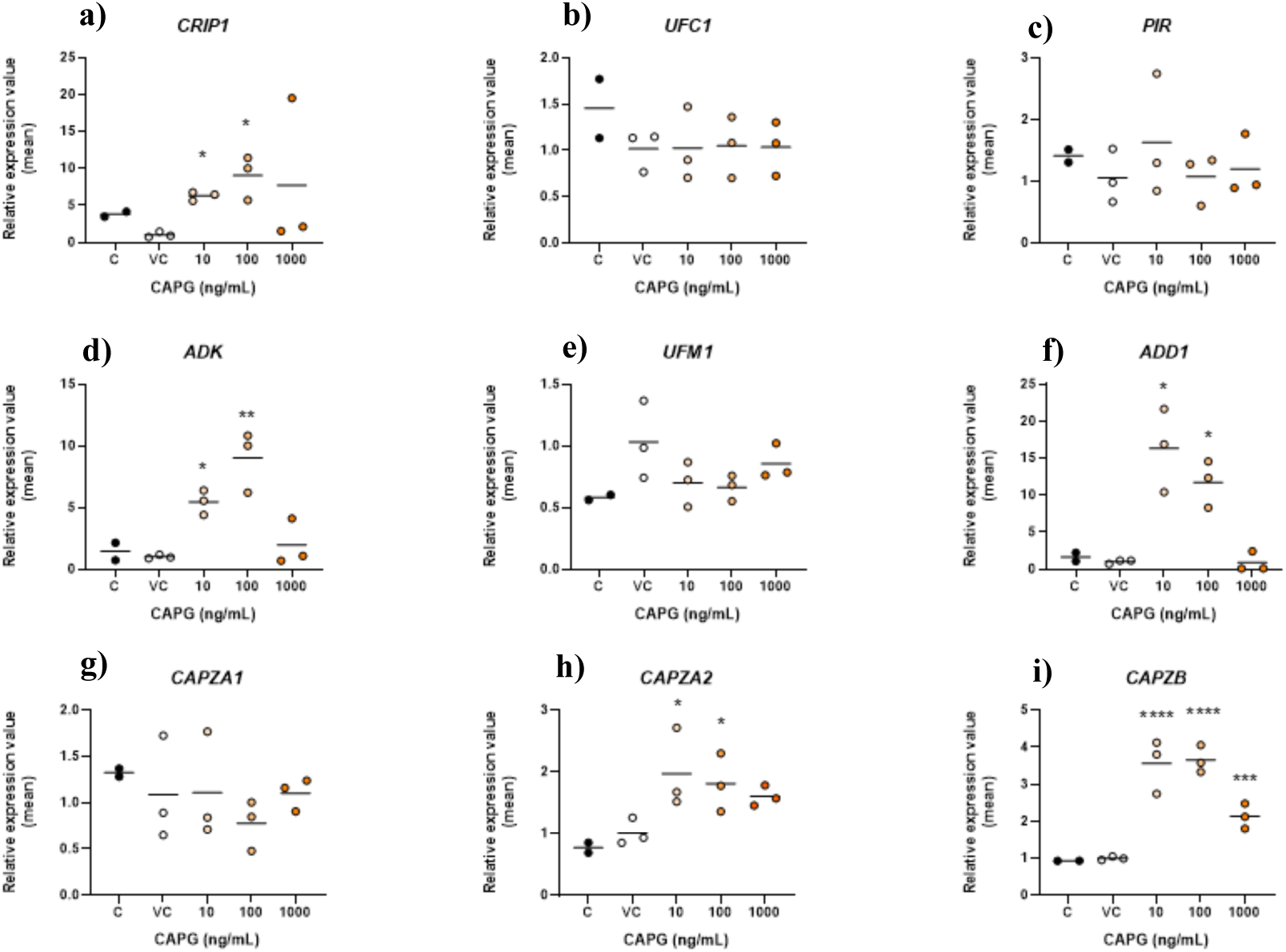
Analysis of expression values by qRT-PCR for selected transcripts in human endometrial epithelial cells (Ishikawa cells: n=3 biological replicates) for **a)** *CRIP1*, **b)** *UFC1*, **c)** *PIR*, **d)** *ADK*, **e)** *UFM1*, **f)** *ADD1*, **g)** *CAPZA1*, **h)** *CAPZA2*, and **i)** *CAPZB*. Cells were treated for 24 hours with i) control (black circles), ii) vehicle control (open circles), iii) 10 ng/ml, iv) 100 ng/ml, or v) 1000 ng/ml (orange circles) recombinant bovine CAPG. Expression values with bar representing mean (calculated with 2^-ΔΔCt^ method using *ACTB, GADPH* and *PPIA* as normaliser transcripts). Significant differences in expression values were calculated by ANOVA analysis using Prism, where samples were compared with a vehicle control of PBS (P<0.05 *, P<0.01 **, P<0.001 ***, P<0.0001 ****).

**Figure 5.**
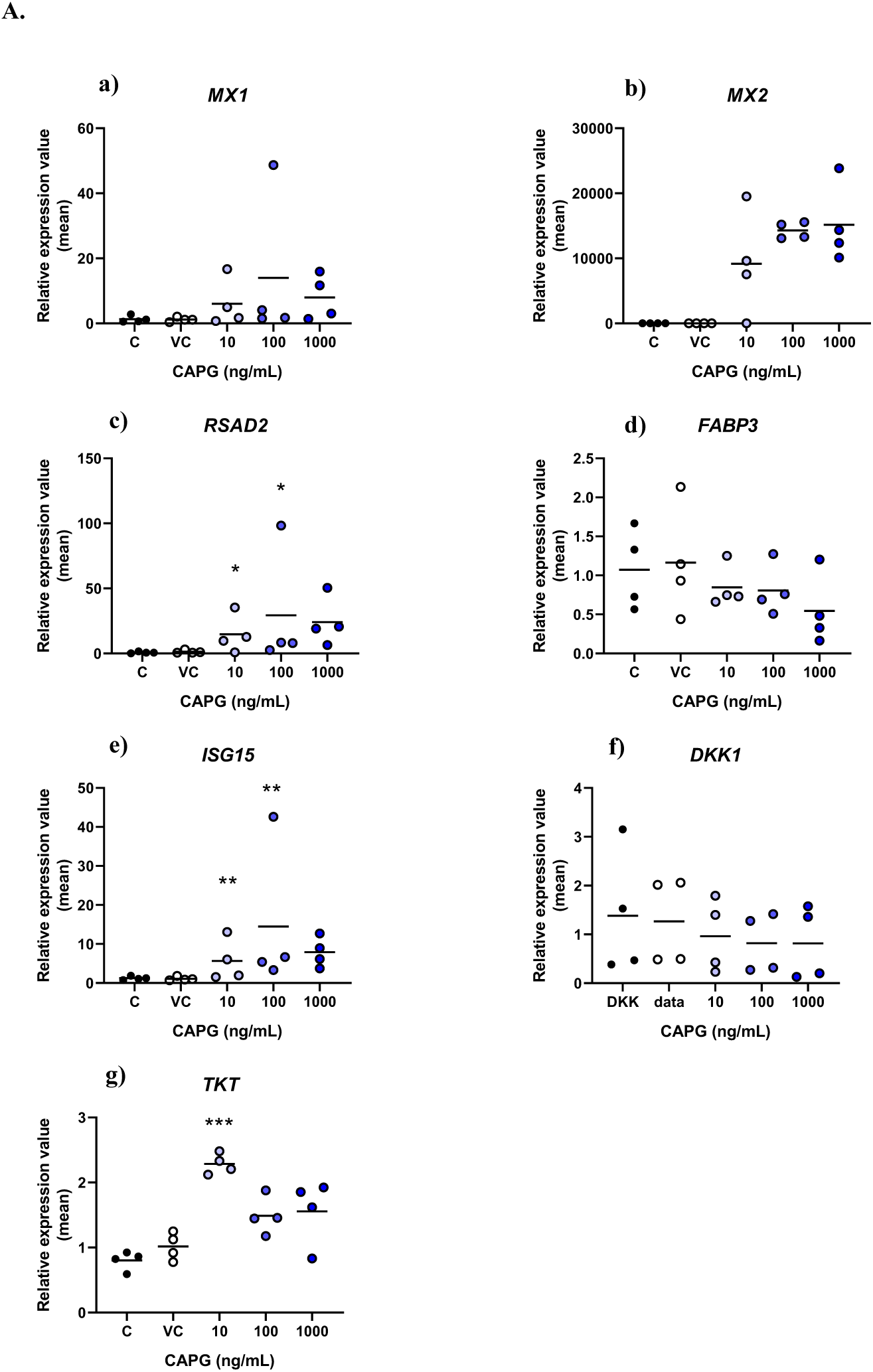
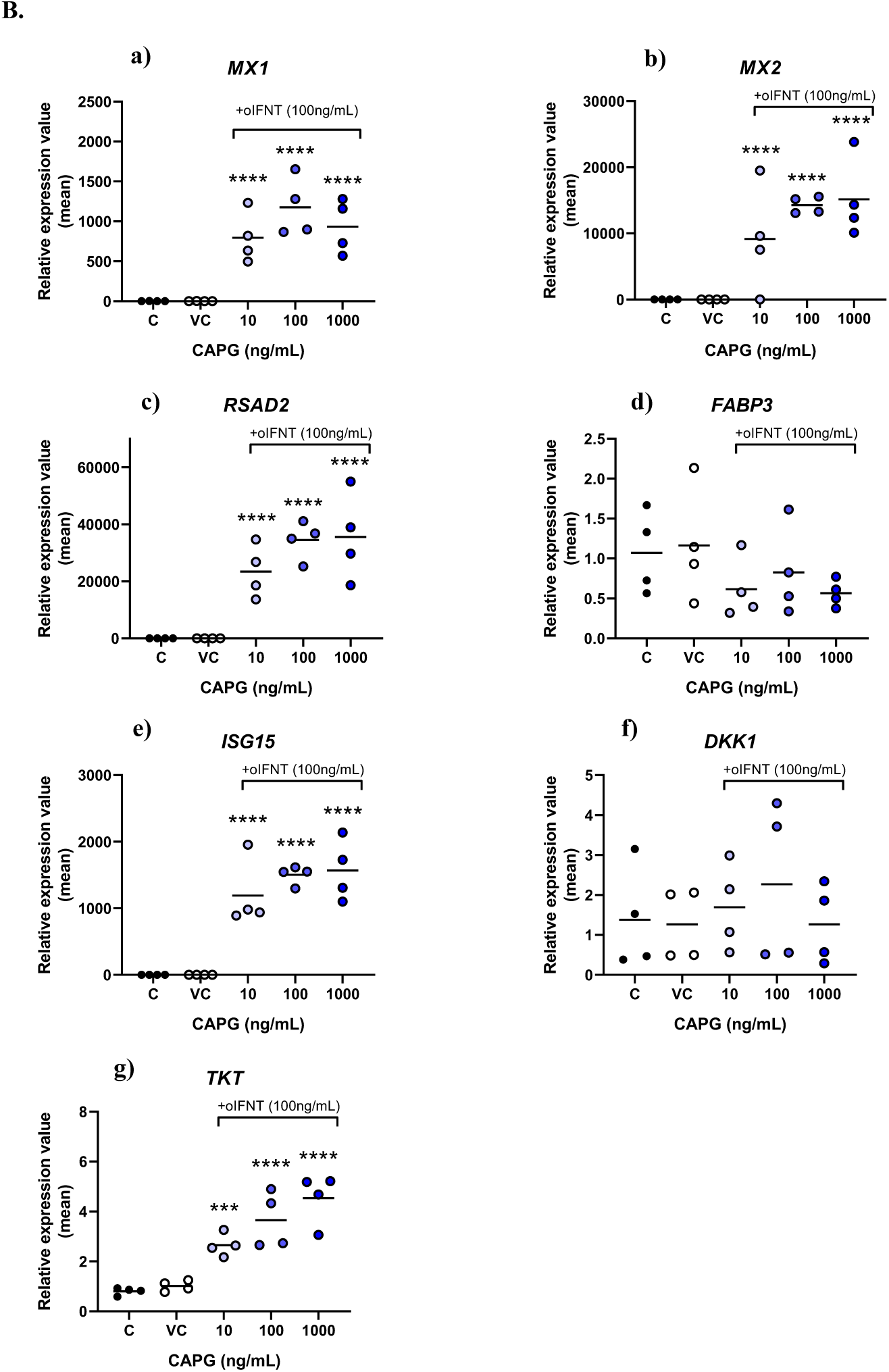
Analysis of expression values by qRT-PCR for selected transcripts in primary bovine endometrial stromal cells (late luteal phase: n=4 biological replicates) for **a)** *MX1*, **b)** *MX2*, **c)** *RSAD2*, **d)** *FABP3*, **e)** *ISG15*, **f)** *DKK1*, **g)** *TKT*, in cells treated for 24 hours with **A)** i) control (black circles), ii) vehicle control (open circles), iii) 10 ng/ml, iv) 100 ng/ml, or v) 1000 ng/ml recombinant bovine CAPG (blue circles). **B)** i) control (black circles), ii) vehicle control (open circles), iii) 10 ng/ml, iv) 100 ng/ml, or v) 1000 ng/ml recombinant bovine CAPG and recombinant bovine IFNT (100 ng/ml) (blue circles). Expression values with bar representing mean (calculated with 2^-ΔΔCt^ method using *ACTB* and *GADPH* as normaliser transcripts). Significant differences in expression values were calculated by ANOVA analysis using Prism, where samples were compared with a vehicle control of PBS (P<0.05 *, P<0.01 **, P<0.001 ***, P<0.0001 ****).

## DISCUSSION

We tested the hypothesis that CAPG that is highly conserved in placental mammals and would facilitate establishing uterine receptivity/ enhance the pregnancy recognition process in species with different implantation strategies, specifically human and cow. Moreover, we hypothesised that high sequence conservation of this protein would be indicative of a function in early pregnancy/ implantation in species with different implantation strategies. We show here for the first time that the CAPG protein coding sequence is highly conserved in the sample of 14 placental mammals investigated. Treatment with a recombinant form of bovine CAPG (produced specifically for this study) modifies the transcriptome of both human and bovine endometrial cells in a species- and cell-specific manner *in vitro*. In cattle, CAPG not only modifies the transcriptional response when present alone, but it also works synergistically with IFNT to enhance the pregnancy recognition process. Addition of recombinant bovine CAPG to human endometrial epithelial cells can also elicit a change in selected transcripts with which it is known to interact in other non-reproductive/implantation physiological systems. We propose that CAPG, a protein with a high degree of sequence conservation across eutherian mammals, plays an integral role in modifying the uterine endometrium in bovine and humans *in vitro*, both of which have different early pregnancy morphologies.

### CAPG protein sequence is highly conserved amongst eutherian mammals

CAPG is a member of the gelsolin family of proteins all of which are comprised of gelsolin-like repeats in their protein sequences, of which CAPG has three (Archer *et al.*, 2004). It is a calcium sensitive protein that reversibly blocks the barbed ends of actin filaments and is predominantly expressed in the cytoplasm of macrophages (Dabiri *et al.*, 1992). CAPG null mice appear to be fertile, but have impaired motility function in macrophages derived from bone marrow (Witke *et al.*, 2001). There is evidence for a duplication event within the gelsolin family of proteins leading to some members of the family having 6 gelsolin subunits, rather than the three that are present in CAPG. It is thought that the family underwent this duplication event in the vertebrate lineage which may explain the sequence similarity across members of the family (Kwiatkowski, 1999). While CAPG null mice are fertile, Campbell *et al.*, (2002) demonstrated that knockout of a close family member (flightless 1: Fliih) resulted in embryo development but subsequent post-implantation loss. Interestingly, if the human form of CAPG was introduced, not only were mouse cells capable of producing the human form, it rescued the post-implantation embryo loss that had been observed in the mice (Campbell *et al.*, 2002). These data suggest that members of this family have a high degree of sequence similarity and family members may play a role in compensatory mechanisms in certain knockout mouse models. Moreover, the data presented here demonstrate that CAPG - with its high level of sequence conservation across species – may play a similar role across eutherian mammals with different implantation strategies.

### CAPG induces a transcriptional response in bovine endometrial epithelial cells that is distinct from that induced by IFNT and may facilitate the pregnancy recognition process

Our data support the hypothesis that CAPG facilitates the pregnancy recognition process working in synergy with IFNT but also by modifying selected transcripts in its own right. Previous studies have demonstrated that there are conceptus-induced changes in the bovine endometrium that occur coordinate with IFNT production but are independent of the actions of IFNT exposure alone. For example, production of prostaglandins by the elongating conceptus pre-primes the endometrium prior to the actions of IFNT (Spencer *et al.*, 2013). Other studies have shown that there are early IFNT-independent changes to the endometrium (Forde, Carter, *et al.*, 2011; Mathew *et al.*, 2019), as well as during the pregnancy recognition period (Bauersachs *et al.*, 2012). Of these transcripts that were identified as conceptus-specific but independent of IFN exposure, we have identified that CAPG modifies 5 of these. The discrepancy may be due to the actions of additional conceptus-derived proteins. An additional >1000 proteins were identified in conceptus-conditioned medium have been identified as produced on Day 16 while 85 proteins are present in the uterine luminal fluid in only pregnant animals on day 16 (Forde *et al.*, 2015).

Treatment of bovine stromal cells with CAPG revealed few differences in gene expression. This may reflect the *in vivo* scenario whereby conceptus-derived products first come into contact with the endometrial epithelium and it is possible that any actions of CAPG that may occur *in vivo* may be indirect via actions on epithelial cells which may produce factors that can themselves act on stromal cells. Unlike the opposite mechanism of the action of stromal-derived growth factors such as HGF (Bazer *et al.*, 2008). These distinct effects on epithelial versus stromal cells reflect the different roles these cells play in facilitating uterine receptivity to implantation.

### High levels of sequence identity between bovine and human CAPG - bovine CAPG can elicit a response in human endometrial epithelial cells but only at low concentrations in vitro

RNA sequencing showed no effect of bovine CAPG on human endometrial epithelial cells. However, sequencing was only carried out on those cells exposed to 1000 ng/ml of CAPG alongside appropriate controls. Selected transcripts known to interact with CAPG were identified from string interaction networks, and qRT-PCR was carried out showing a dose response of human cells to recombinant bovine CAPG but only at the lower concentrations, *i.e.* 10 and 100 ng/ml (Figure 4), except for the transcript *CAPZA1*. The transcripts that were altered by CAPG in human endometrial epithelial cells are those that have been previously identified as interacting with CAPG in different systems in humans, thus implicating CAPG in modifying endometrial transcripts in the human endometrium. The lack of DEGs in human cells treated with the highest concentration of rbCAPG may simply reflect a dose-specific mechanism of action in humans. Indeed, no data are available to date that demonstrate the concentration of CAPG present in the uterine lumen in humans. CAPG is expressed in the human embryo (Stirparo *et al.*, 2018) as well as in the embryos of other species including bovine (Nakamura *et al.*, 2016) and porcine (Whitworth *et al.*, 2011) further implicating CAPG in conceptus-maternal interactions in a range of mammals.

In conclusion, we have demonstrated that CAPG, a protein that is highly conserved in sequence across different eutherian mammals, modifies both mRNA and miRNA in endometrial epithelial cells in a species-specific manner. In cattle, it modifies transcripts important for pregnancy recognition, but also works in synergy with IFNT to facilitate the process of pregnancy recognition. In humans, it modifies specific interaction partners but only at lower concentrations *in vitro*. CAPG modifies miRNAs in the endometrial epithelia of both humans and cattle but does so in a species-specific manner. We propose that CAPG plays a facilitative role in establishing uterine receptivity to implantation and contributes to early pregnancy success across species with different early implantation strategies.

## ACKNOWLEDGEMENTS

We would like to Dr Fuller Bazer from Texas A&M for the kind gift of recombinant ovine IFNT. We acknowledge Dr James Cronin from the University of Swansea for guidance with the bovine cell isolation method. We would like to acknowledge the assistance of Stefania Mountevedi in helping with the bovine cell isolation and culture. We would like to thank Dr Ian Carr, Morag Raynor and Ummey Hany from the University of Leeds’s next generation sequencing facility core for undertaking sequencing analyses. This work was supported by N8 agri-food pump priming, QR GCRF, as well as BBSRC grant number BB/R017522/1.

